# Newly identified proviruses in Thermotogota suggest that viruses are the vehicles on the highways of interphylum gene sharing

**DOI:** 10.1101/2020.11.07.368316

**Authors:** Thomas H. A. Haverkamp, Julien Lossouarn, Olga Zhaxybayeva, Jie Lyu, Nadège Bienvenu, Claire Geslin, Camilla L. Nesbø

## Abstract

Phylogenomic analyses of bacteria from the phylum Thermotogota have shown extensive lateral gene transfer (LGT) with distantly related organisms, particularly with Firmicutes. One likely mechanism of such DNA transfer is viruses. However, to date only three temperate viruses have been characterized in this phylum, all infecting bacteria from the *Marinitoga* genus. Here we report 17 proviruses integrated into genomes of eight Thermotogota genera and induce viral particle production from one of the proviruses. The proviruses fall into two groups based on sequence similarity, gene synteny and taxonomic classification. Proviruses of one group are found in six genera and are similar to the previously identified *Marinitoga* viruses, while proviruses from the second group are only distantly related to the proviruses of the first group, have different genome organization and are found in only two genera. Both groups are closely related to Firmicutes in genomic and phylogenetic analyses, and one of the groups show evidence of very recent LGT and are therefore likely capable of infecting cells from both phyla. We conjecture that viruses are responsible for a large portion of the observed gene flow between Firmicutes and Thermotogota.

## Introduction

The phylum Thermotogota comprises anaerobic fermentative bacteria, most of which are thermophiles [1]. They are common in subsurface environments such as marine vents, terrestrial hot springs and deep subsurface oil reservoirs [2–5]. On phylogenetic trees of 16S rRNA gene, Thermotogota are usually a deep branching bacterial lineage, while ribosomal proteins and other markers do not always agree with that placement [6, 7]. Such discrepancies are likely due to lateral gene transfer (LGT), which has been an important force shaping the genomes of Thermotogota, with Firmicutes and Archaea being their most notable gene transfer partners [1, 7, 8]. The LGT between Firmicutes and Thermotogota is so extensive that the two phyla have been suggested to be linked by ‘‘highways of gene sharing’’ [7]. However, how these inter-phylum gene-sharing events occur is still unclear.

The subsurface constitutes the largest biosphere on Earth and is estimated to contain ~70% of all cells [9]. Viruses are likely to be particularly important in subsurface environments, since 97% of all viruses on earth being found in soil and sediments [10, 11]. Moreover, although both prokaryotic cell and virus numbers decrease with depth, the virus-to-cell ratio increases with depth [10, 12, 13]. Phylogeographic studies of hyperthermophilic *Thermotoga* and mesophilic *Mesotoga* have revealed genetic interaction between geographically distant populations, particularly among the hyperthermophilic *Thermotoga* [3, 5]. Viruses are one potential source of such long-distance dispersal of genetic material [14], especially for anaerobic organisms where surface dispersal is problematic.

Although viruses are likely candidates for transferring DNA both within and between species, only three temperate siphoviruses (MCV1, MCV2, and MPV1), all infecting one Thermotogota genus, *Marinitoga*, have been described [15, 16]. MCV1 and MCV2 infect *Marinitoga camini* strains isolated from deep-sea hydrothermal vents [16]. MPV1 infects the deep-sea marine vent bacterium *Marinitoga piezophila*, where it is highjacked by a plasmid co-occurring in the same host, illustrating the potential route of gene mobilization in these ecosystems [15]. The three viruses are found as proviruses in their host genomes and show similar genomic organization and virion morphology. Phylogenetic and protein sequence-similarity analyses of the viral ORFs revealed that they often group either with Firmicutes or Firmicutes’ viruses, which suggests that viruses infecting members of Firmicutes and Thermotogota phyla share a common gene pool [15, 16].

Here we report 17 additional proviruses in Thermotogota genomes from eight Thermotogota genera, and a successful induction of one of these proviruses. The identified proviruses fall into two distinct groups. Both groups are closely related to Firmicutes viruses, and the proviruses from one of these groups are likely able to infect cells from both phyla. We hypothesize that membrane transport proteins, such as ABC transporters, serve as receptors for Thermotogota viruses. We propose a mechanism that could account for the highways of gene sharing observed between Thermotogota and Firmicutes, where LGT of viral genes encoding transmembrane proteins may make the host vulnerable to new viruses.

## Material and Methods

### Prediction and taxonomic classification of proviruses and functional annotation of their ORFs

One hundred eleven Thermotogota genomes were downloaded from either Genbank or IMG [17] prior to June 2018. For draft genomes, the contigs were combined into ‘artificially closed’ genome using the “union” command from the EMBOSS package (version 6.6.0) [18]. Each genome was screened for the presence of proviruses using the Prophinder web server [19], PHAST web server [20], and PhiSpy (version 2.3) [21] between October 2014 and July 2018 (**Supplementary Table S1**). For artificially closed genomes, the proviral regions that crossed contig borders were discarded. Putative provirus regions were inspected to identify the most likely provirus sequence by 1: looking at annotations, 2: identifying possible flanking tRNA genes and 3: comparing the region to genomes from the same genus and defining the boundaries, to ensure that flanking genomic regions present in closely related genomes without provirus were not included. Proviruses were considered complete if they contained modules for lysogeny, replication, packaging, head/tail morphogenesis and lysis. If one of these modules were missing, the provirus was scored as incomplete.

To see if there were close relatives of the predicted proviruses in other genomes, provirus ORFs were used as queries in BLASTX (ver.2.2.26) [22] searches of the NCBI non-redundant (nr) database [23] (accessed between July 2018 and March 2020). When homologs for multiple ORFs from the predicted provirus were found in the same distantly related subject genome (usually a Firmicutes genome), the identified genome was downloaded and aligned to the Thermotogota genome carrying the provirus using Progressive Mauve [24]. This resulted in the identification of a local alignment covering similar proviruses in two otherwise distantly related genomes. The aligned region was used to determine the boundaries of the provirus, limiting the provirus ends to the ends of the alignment.

Proviral-ORF annotations were obtained from their respective Genbank entries and supplemented by results from BLASTP searches [22] of the *nr* database with an expected-value cutoff of 10^−1^, and from HHpred searches [25] of the PDB database [26] with a probability cutoff of 99%. In addition, recombinase- and terminase-encoding ORFs were annotated using InterProScan [26], as implemented in Geneious v.10 (Biomatters Ltd.).

The sequences of the predicted proviruses were compared to each other using BLASTN and TBLASTX (ver.2.2.26) [22] and visualized using genoPlotR [27] and Circos [28]. Taxonomic classification of the provirus genomes was carried out using searches of NCBI’s viral RefSeq database (v. 94) as implemented in VContact2 using Diamond [29] to identify viral protein clusters and ClusterONE [30] to obtain virus clusters [31, 32]. Taxonomic classification was also assessed with Virfam [33], VIRIDIC [34] and VIPTree [35]. Morphological classification was obtained with Virfam [33].

#### Inference of potential host range of the putative Thermotogota viruses

A database containing all proteins from 59 Thermotogota genomes without identified proviruses was constructed in Geneious v.10. Translated Thermotogota provirus proteins were used as queries in BLASTP searches of this database. The provirus genes were scored as present in the Thermotogota genomes if the query protein had a match with > 50% amino acid identity and > 60% coverage.

CRPISPR spacer sequences from 90 Thermotogota genomes were obtained from IMG [17]. The spacers were mapped to the provirus genomes in Geneious v.10, allowing upto 10% nucleotide mismatches.

#### Phylogenetic analyses of provirus genes and candidate receptor genes

Homologs of provirus genes selected for phylogenetic analysis were obtained by searching each translated proviral gene against *nr* database (accessed between December 2019 and March 2020), as well as a local database of all the *Thermotogota* virus proteins identified, using BLASTP (version 2.2.26), with E-value cutoff of 10^−1^. The 20 top-scoring matches from each database were retrieved and aligned using MAFFT v. 7.450 with the G-INS-I option [36]. Identical sequences and highly similar sequences from the same genus were removed. Alignment positions with > 50% gaps were trimmed. Phylogenetic trees were reconstructed using RAxML [37] with WAG+G substitution model with four rate categories and 100 bootstrap replicates, as implemented in Geneious v.10.

Candidate receptor proteins in genomes of *Petrotoga* sp. 8T1HF07.NaAc.6.1, *Petrotoga olearia*, *Petrotoga mobilis*, *Petrotoga* sp. 9T1HF07.CasAA.8.2, *Defluviitoga tunisiensis*, *Lacticigenium naphtae*, and *Mahella australiensis*, which had proviruses assigned to Group 2 (see the Results section for definition), were identified in IMG using an amino acid identity cut-off of 50%. Homologs in *Geosporobacter ferrireducens* genome, which was not available in IMG, were identified using BLASTP search with E-value cutoff of 10^−10^. Collection of additional homologs and phylogenetic analyses were carried out as described above.

Phylogenetic analysis of single copy gene in Thermotogota genomes available in Genbank (accessed May 27 2020) were done using the GToTree pipeline [38] with the Bacterial hmm-set of 74 target genes. The resulting alignment was imported into Geneious Prime 2020.1.2 where sites with more than 50% gaps were removed, giving an alignment of 11,003 amino acid positions. The phylogenetic tree was reconstructed using FastTree with the JTT model with optimized Gamma20 likelihood [39].

#### Virus induction and electron microscopy

*T. africanus* H17ap6033 and two *Petrotoga* isolates, *P.olearia* and *Petrotoga* sp. 8T1HF07.NaAc.6.1, were cultivated in a modified Ravot medium as previously described [15] at 65°C and 55°C, respectively. Attempts were made to increase the viral production of the strains by using mitomycin C, as reported previously [15, 16]. A final concentration of 5 μg/mL of mitomycin C was added to 300 mL bacterial culture at early to mid-log growth phase. After 3 hours of incubation with mitomycin C, cultures were centrifuged at 7500 rpm and 4°C for 15 min, and supernatants were ultracentrifuged at 37 000 rpm (~100 000 g) and 10°C for 1h (Beckman Optima LE-80 K; rotor 70.1.Ti). Pellets were resuspended in 100μL of buffer (10 mM Tris-HCL, 100 mM NaCl, 5 mM CaCl_2_, 20 mM MgCl_2_) and suspensions were prepared for negative staining electron microscopy as previously described [40]. Briefly, 5 μL of the suspensions were directly spotted onto a Formwar carbon coated copper grid. Putative virus-like particles were allowed to adsorb to the carbon layer for 2 min and excess of liquid was removed. 5 μL of a staining uranyl acetate solution (2%) was then spotted to the grid for 45 s and excess of liquid was removed again. The grid was imaged at 120 kV in a JEOL JEM 100 CXIIVR transmission electron microscope.

## Results

### Newly identified Thermotogota proviruses come from two distinct viral lineages

Analysis of 111 Thermotogota genomes identified 20 proviruses, including the three already characterized viruses from *Marinitoga* [15, 16] and four likely partial proviruses (**Supplementary Table S1** and **Supplementary Table S2**). One of the 20 proviruses is present with 100% nucleotide sequence identity in all six available *Thermosipho melanesiensis* genomes [41], and therefore is counted as just one novel provirus. An additional provirus (MLaV1) was reported in a *Marinitoga lauensis* genome after we completed the screening [42]. Due to its similarity to proviruses identified in other *Marinitoga* genomes, it was not included in our further analyses.

The predicted proviruses can be divided into two distinct groups, hereafter denoted as Group 1 (15 proviruses: 13 complete and 2 incomplete) and Group 2 (5 proviruses: 3 complete and 2 incomplete). First, the genome organization differs between the proviruses in two groups (**Fig. 1** and **Fig. 2**). Second, the genes within each group are more similar than the genes between groups (**Fig. 3, Supplementary Table S2**). Third, the two groups form separate clusters in the VContact2 network (**Supplementary Fig. S1, panel A**). Finally, the two groups show up as different clades on the Viral Proteomic Tree (**Supplementary Fig. S2**). Group 1 proviruses are found in the genera *Marinitoga*, *Thermosipho*, *Kosmotoga*, *Mesotoga*, *Geotoga* and *Mesoaciditoga*, while Group 2 proviruses are limited to the genera *Petrotoga* and *Defluviitoga* (**Supplementary Fig. S1, panel B**). However, presence of 29 protein families shared between the two groups **(Supplementary Table S3)**suggests that LGT may occur between the viruses of the two groups.

**Fig. 1.**
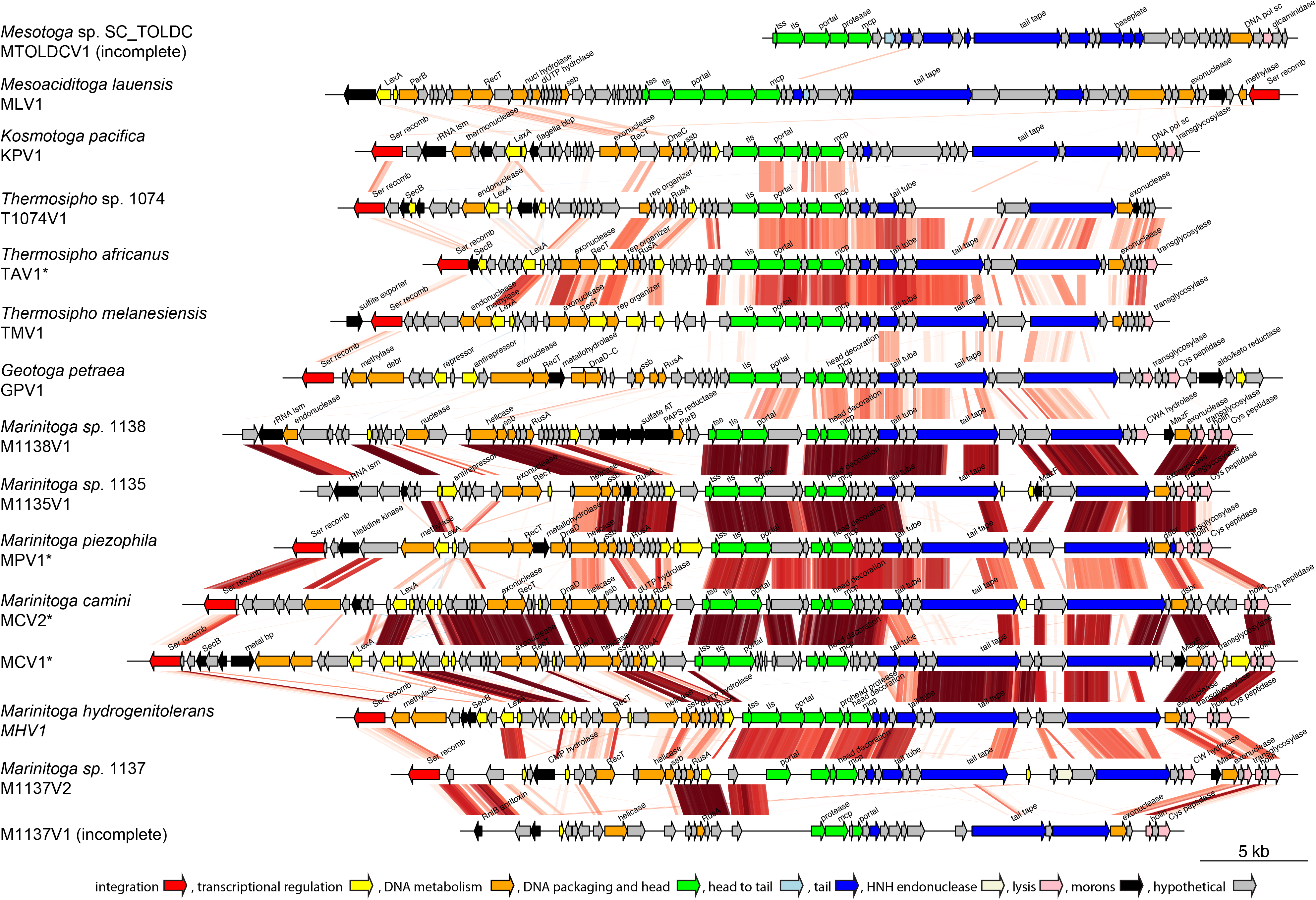
Comparison of sequences from all detected Group 1 proviruses. Provirus name and the species of its host are shown to the left of the nucleotide sequence, in which predicted ORFs are depicted as arrows. Proviruses that have been induced and shown to produce virus particles are marked with an asterisk. The lines connect regions of adjacent viruses that have TBLASTX similarity of more than 30% over 100bp. Lines are colored in red or blue indicate that the matching sequences encoded in the same or opposite strand, respectively. The predicted ORFs are color-coded based on their function and should be considered approximate, because it relies only on gene annotations. Selected gene annotations are included and abbreviated as follows. Ser recomb: serine recombinase, LexA: LexA repressor, ParB: ParB-like nuclease, RecT: RecT family recombinase, dUTP hydrolase: deoxyuridine 5’-triphosphate nucleotidohydrolase, ssb: single stranded DNA-binding protein, tss: terminase small subunit, tls: terminase large subunit, mcp: major capsid protein, tail tape: tail tape measure protein, rRNA lsm: ribosomal RNA large subunit methyltransferase, flagella bbp: flagella basal-body protein, DnaC: DnaC replication protein, DnaD: DnaD replication protein, DNA pol sc: DNA polymerase sliding clamp, SecB: SecB protein-export protein, rep organizer:replisome organizer, RusA: RusA family crossover junction endodeoxyribonuclease, Cys peptidase: cysteine peptidase, sulfate AT: sulfate adenylyltransferase subunit 2, PAPS reductase: phosphoadenosine phosphosulfate reductase, CW hydrolase: cell wall-associated hydrolase, MazF: MazF endoribonuclease, dsbr: DNA double-strand break repair protein, metal bp: metal-binding protein, CMP hydrolase: cytidine 5′-monophosphate hydrolase. The figure was produced using genoPlotR [27].

**Fig. 2.**
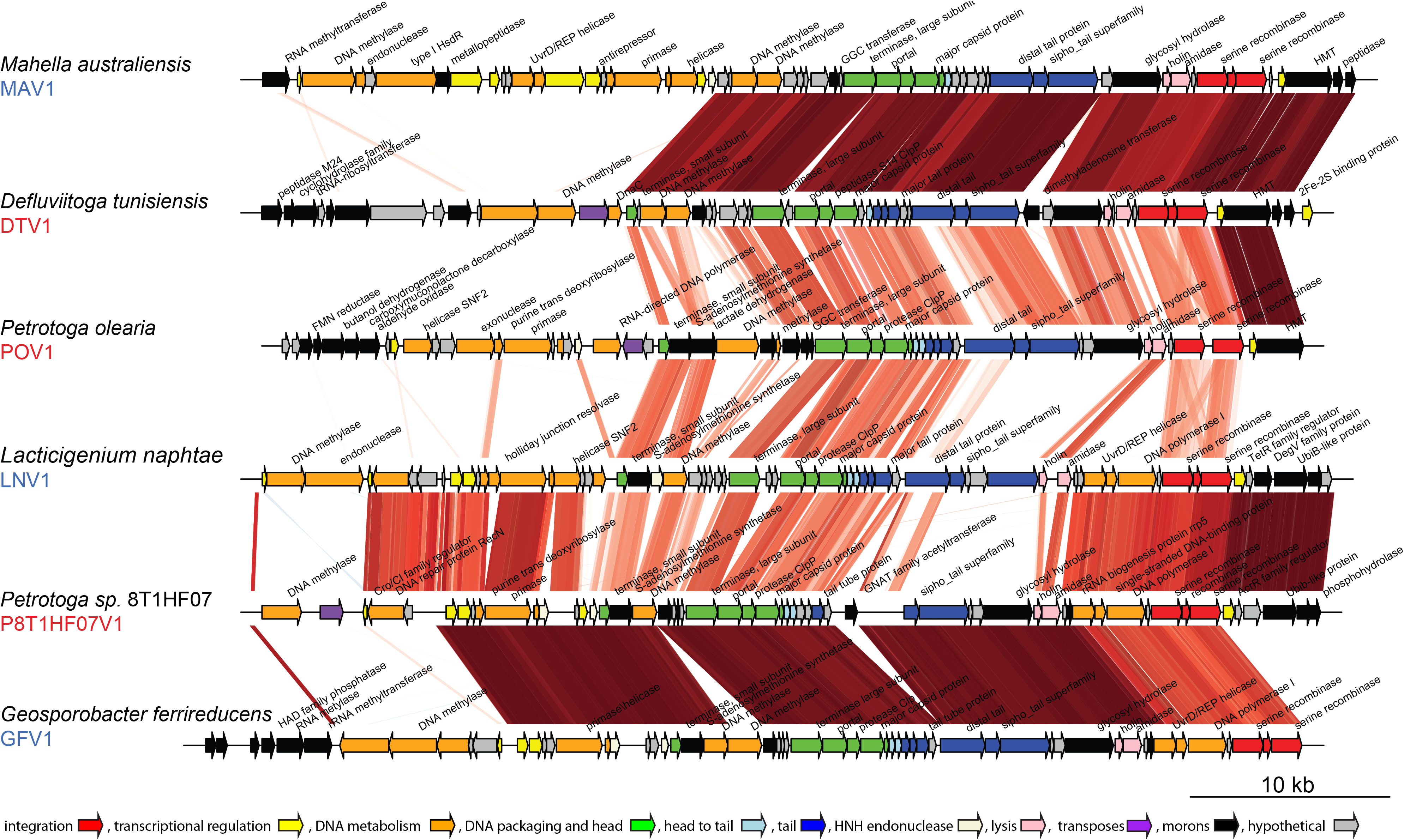
Comparison of sequences from three complete Thermotogota Group 2 proviruses and their Firmicutes’ homologs. Provirus name (in red for Thermotogota and blue for Firmicutes) and the species of its host are shown to the left of the nucleotide sequence, in which predicted ORFs are depicted as arrows. The lines connect regions of adjacent viruses that have TBLASTX similarity of more than 30% over 100bp. Lines are colored in red or blue indicate that the matching sequences encoded in the same or opposite strand, respectively. The predicted ORFs are color-coded based on their function and should be considered approximate, because it relies only on gene annotations. Selected gene annotations are included and abbreviated; HMT ATPase: heavy metal translocating ATPase, FMN reductase: flavine mono nucleotide reductase, HAD family phosphatase: haloacid dehalogenase superfamily of hydrolase). The figure was produced using genoPlotR [27].

**Fig. 3.**
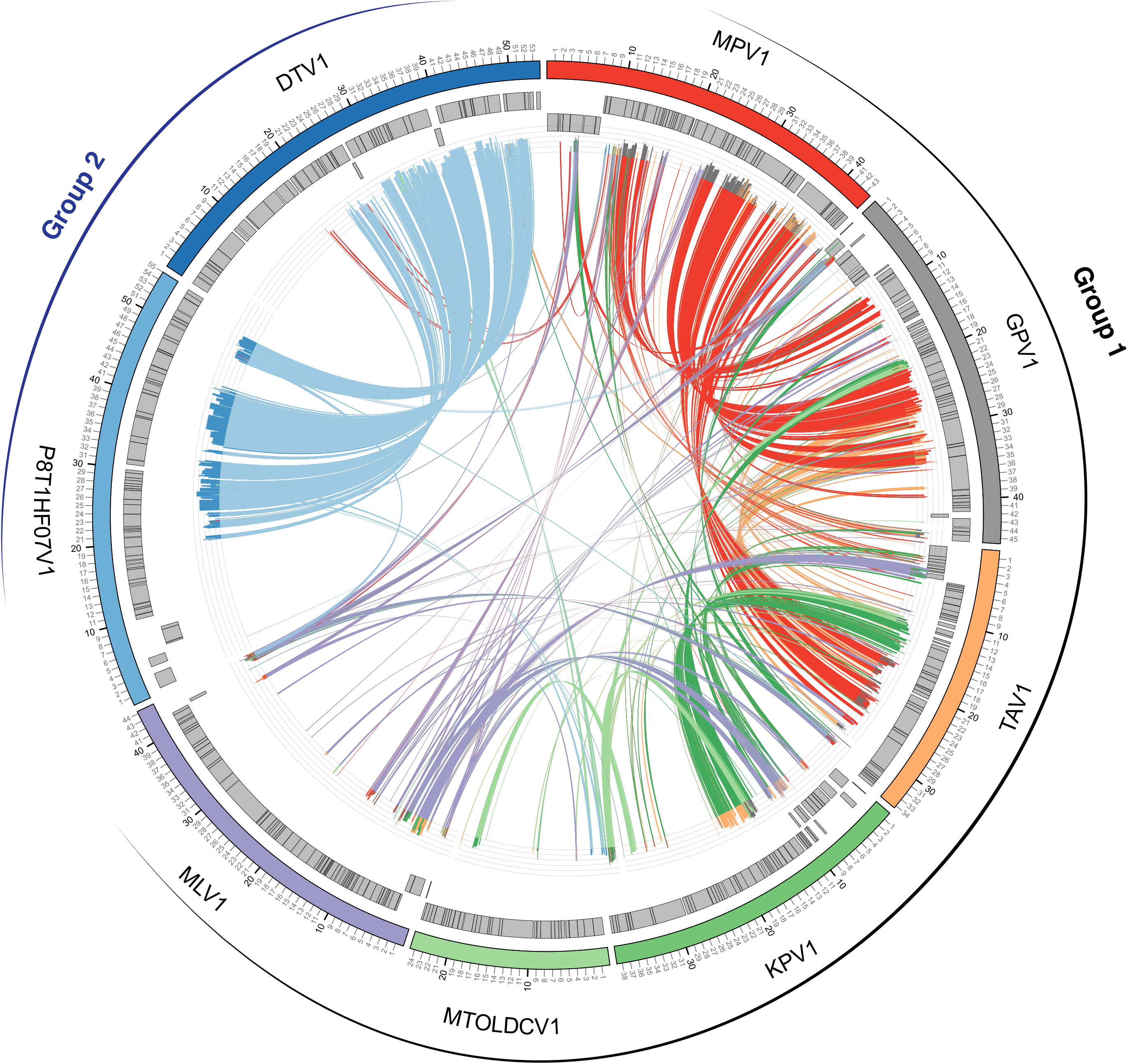
Comparison of representative Thermotogota proviruses. Due to sequence similarity, only one provirus per Thermotogota genus is shown. The nucleotide sequences of the proviruses are arranged around the circle and color-coded. Numbers indicate kilobases (kb) and grey boxes outline locations of predicted genes. Lines connecting different proviral sequences represent TBLASTX matches between the proviral regions, with the percent identity shown in histograms at the ends of each line. The plot was created using Circos [28].

### Classification and genomic features of Group 1 proviruses

All complete Group 1 proviruses are likely to encode siphoviruses based on their head, neck and tail gene sequences [33], and the morphology observed for the earlier characterized MPV1, MCV1, MCV2 viruses and the TAV1 virus induced in the current study (see below).

None of the proviruses have significant nucleotide identity with viral genomes in the NCBI nr/nt database. Following taxonomic criteria, where viruses with > 50-70% nucleotide identity over the full genome belong to the same genus and viruses with > 95% nucleotide identity belong to the same species [43, 44], the complete proviruses are assigned to 13 new species and at least 11 new genera (**Supplementary Table S4**). The sequence similarity suggests that the closest relatives of the Group 1 proviruses are Firmicutes’ viruses (**Supplementary Fig. S1, panel A)**, since 15% of the provirus genes have Firmicutes as the top-scoring match, if members of the proviruses’ host genus are excluded (**Supplementary Table S5**).

The proviruses have the same modular structure as genomes of the earlier described MPV1, MCV1 and MCV2 viruses [15, 16] (**Fig. 1**). The 5’ module contains genes involved in lysogeny and is encoded on the opposite strand compared to the rest of the virus genes. The lysogeny module is followed by modules for replication, packaging, morphogenesis and host lysis. Similar to the described *Marinitoga* viruses [15, 16], the gene content of lysogeny module of all examined proviruses is very variable, with only the recombinase gene conserved (**Fig. 1**).

All Group 1 proviruses are inserted next to a tRNA gene. Eight of them (the *Marinitoga* proviruses MPV1, MCV1, MCV2, MHV1 and M1137V2; the *Kosmotoga pacifica* provirus KPV1, *Geotoga petrae* provirus GPV1, and *Mesoaciditoga lauensis* provirus MLV1), are inserted next to the tRNA-Glu gene, and carry similar site-specific DNA serine recombinases (KEGG Orthology; K06400, homologs of Marpi_0291 in MPV1 from *M. piezophila*) (**Fig. 4**). The *Thermosipho* proviruses TMV1 and T1074V1 carry more distant homologs of this recombinase (**Fig. 4**) and are inserted next to the tRNA-Phe gene. The most divergent homolog of the serine recombinase is present in TAV1, which is inserted next to the tRNA-Pro gene. MTOLDCV1 is also inserted next to a tRNA-Pro gene, but it is located at the end of a contig in an incomplete single cell genome and its 5’ end (where recombinase would be found) is missing (the *Mesotoga* sp. SC_TOLDC recombinase included in **Fig. 4** is located on a separate contig). We hypothesize that these recombinases are integrases that specifically recognize the tRNA genes next to which the provirus is inserted (i.e., the tRNA-Glu, tRNA-Phe, or tRNA-Pro genes). The remaining three proviruses (M1135V1, M1138V1, and M1137V1) are inserted next to tRNA-Cys gene and may also use similar integration mechanism, but these proviruses do not have the detectable serine recombinase homologs. M1135V1 and M1138V1 have homologous ORFs of unknown function in the recombinase gene position (**Fig. 1**), which may or may not provide this function.

**Fig. 4.**
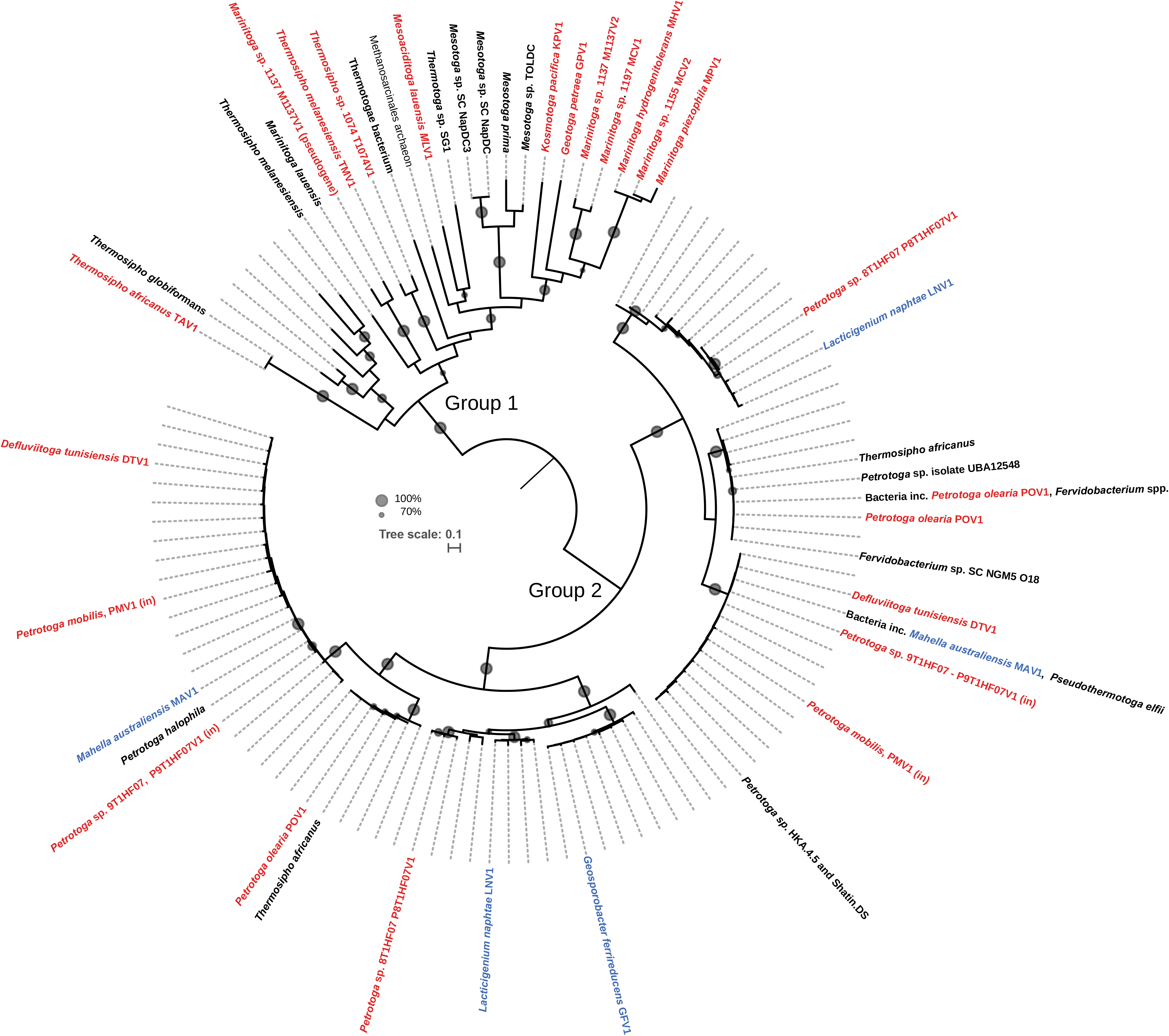
Maximum likelihood tree of recombinases found in Thermotogota proviruses and of their homologs in Firmicutes proviruses, and Thermotogota and Firmicutes genomes. Host names of Thermotogota and Firmicutes proviruses are colored in red and blue, respectively. The names of their proviruses are added next to the host name. Names of Thermotogota homologs that either resided outside of proviral regions or come from a genome without detected proviruses are shown in black. Branches without labels represent Firmicutes without an identified Group 2 provirus. Homologs from incomplete proviruses are labeled with “(in)”. Circles on the branches represent bootstrap support, and only values above 70% are shown. Some proteins have identical amino acid sequences in more than one organism. The protein labelled ‘Bacteria inc. POLV1 *Fervidobacterium* spp.’corresponds to accession number WP_011994748.1 and is found in *Fervidobacterium nodosum* Rt17-B1 (NC_009718.1), *Fervidobacterium pennivorans* DSM 9078 (NC_017095.1), *Fervidobacterium islandicum* (NZ_CP014334.1), *Fervidobacterium gondwane*nse DSM 13020 (FRDJ01), *Petrotoga olearia* (PNR98053) and *Coprothermobacter proteolyticus* (PXJB01). The protein labelled ‘Bacteria inc. *Mahella australiensis* MAV1, *Pseudothermotoga elfii*’ corresponds to accession number WP_013782344.1 and is also found in *Clostridium* sp. SYSU GA15002T (NZ_CP040924.1), *Thermoanaerobacter thermocopriae* JCM 7501 (NZ_KI912455.1), *Pseudothermotoga elfii* and MAV1 from *Mahella australiensis.* The tree was rooted by mid-point rooting and visualized using iTOL [56]. Tree scale, substitutions per site.

Another typical viral protein that shows variation across the Group 1 proviruses is the large subunit of the terminase protein involved in the packaging of viral DNA into the virus particle [45]. The proviruses carry three types of these proteins (BLASTP E-value cut-off < 0.01, identity < 25%, **Supplementary Fig. S3**). The first type, exemplified by the protein in MPV1 (Marpi_0320), contains a PBSX family domain and has homologs in MCV1 and MCV2 and the proviruses in *Marinitoga* sp. 1135, *Marinitoga* sp. 1138, *Thermosipho* sp. 1074 and *M. lauensis*. The second, exemplified by the terminase in TAV1 (H17ap60334_04902), contains a ‘Terminase_lsu_T4-like’ domain and has homologs in the proviruses from *K. pacifica*, *T. melanesiensis*, *Mesotoga* sp. TolDC and *G. petrae*. The third type is found in the proviruses in *M. hydrogenitolerans* and *Marinitoga* sp. 1137 (BUA62_RS02495, LN42_01905 and LN42_00550). These terminases also contain a ‘Terminase_lsu_T4-like’ domain and are distant homologs of the second terminase type.

In addition to the recombinase and terminase, other typical viral proteins such as tail tape measure, capsid and portal proteins were identified, but did not always show detectable similarity among the proviruses (**Fig. 1**). Two transcription regulators (Marpi_0297 and Marpi_0298 in MPV1), a DNA repair exonuclease (Marpi_0340 in MPV1) and a single stranded DNA-binding protein (Marpi_0306 in MPV1), show relatively high (32-100%) identity across most Group 1 proviruses (**Fig. 1, Supplementary Table S4**). Genes encoding two hypothetical proteins (homologs of Marpi_0299 and Marpi_0338 in MPV1) are shared among 10 of the proviruses (36-96% identity), suggesting these genes may provide important viral functions.

### Broad host range of Group 1 proviruses

Detection of Group 1 proviruses in the genera *Marinitoga*, *Thermosipho*, *Kosmotoga*, *Mesotoga*, *Geotoga* and *Mesoaciditoga* (**Supplementary Table S1**), suggests that the Group 1 viruses are widespread among Thermotogota, particularly among organisms inhabiting hydrothermal vents. Such wide distribution and relatively high sequence identity among the proviral genomes (**Fig. 1, Supplementary Table S4**) suggest that the Group 1 temperate viruses might have broad host ranges. Experiments showing that MPV1 from *M. piezophila* can infect and transfer a plasmid to a *Thermosipho* isolate is consistent with this hypothesis [15].

Further support comes from mapping of CRISPR spacer sequences from 90 Thermotogota genomes to the Group 1 proviruses. Five of the 17 proviruses matched CRISPR spacers in the genomes from a different genus (**Table 1**). For example, the *Thermosipho* provirus TAV1 had 35 matches to spacers in the genomes of *Pseudothermotoga* and *Thermotoga* spp. Anecdotal evidence corroborates an ability of TAV1 to infect *Thermotoga* spp. Back in 2005, when the sample from the Hibernia oil reservoir containing TAV1 and its host, *T. africanus* H17ap60334, was being processed by one of us (Camilla L. Nesbø) in the laboratory, *Thermotoga* isolates from Troll oil reservoir in the North Sea, which were at the same time being transferred to fresh media, experienced a mass death. Analysis of the genomes of the surviving *Thermotoga* isolates [3] revealed presence of three CRISPR spacer matching the TAV1 genome (**Supplementary Fig. S4**). These spacers were located in the middle of the CRISPR arrays, indicating that they were not new acquisitions [46]. Therefore the only surviving isolates of *Thermotoga* must have had already experienced and survived TAV1 or related virus infections in the oil reservoir.

**Table 1.** CRISPR spacer matches to provirus genomes in Thermotogota genomes. Matches to spacers from the provirus’ host genome are labeled as a self-match.

| provirus name | Genome with a CRISPR spacer match (number of spacers) |
| --- | --- |
| KPV1 | *Kosmotoga olearia* (1) |
| MHV1 | *Pseudothermotoga elfii* NBRC107921 (1), *Marinitoga* sp. 1154 (1) |
| MCV1* | *Marinitoga* sp. 1154 (2) |
| MCV2* | *Marinitoga* sp. 1155 (1) self-match, *Marinitoga* sp. 1154 (2) |
| MPV1* | *Marinitoga* sp. 1137 (1) |
| M1135 | *Marinitoga* sp. 1155 (1) |
| **M1137V1** | *Pseudothermotoga elfii* NBRC107921 (1), *Marinitoga* sp. 1135 (2) |
| **M1137V2** | *Marinitoga piezophila* (1) |
| **M1138** | *Thermosipho africanus* TCF52B (1),<br>*Thermosipho melanesiensis* (1), *Pseudothermotoga elfii*<br>NBRC107921 (1), *Marinitoga* sp. 1137 (1) |
| **TAV1\*** | *Thermosipho africanus* H17ap60334 (3) self -match, *Thermosipho africanus* TCF52B (2), *Thermosipho africanus* Ob7 (1), *Thermosipho melanesiensis* (2), *Thermotoga maritima* 2812B (1), *Thermotoga* sp. EMP (1), *Thermotoga* sp. XYL54 (3), *Thermotoga* sp. CELL2 (3), *Thermotoga* sp. TBGT1766 (3), *Thermotoga* sp. TBGT1765 (4), *Thermotoga* sp. A7A (1), *Thermotoga* sp. MC24 (2), *Pseudothermotoga elfii* lettingae (4), *Pseudothermotoga elfii* NBRC107921 (9), *Pseudothermotoga elfii* DSM9442 (2) |
| **TMV1** | *Thermosipho melanesiensis* (1) self-match, *Thermosipho africanus* TCF52B (1), *Thermosipho africanus* Ob7 (1), *Thermotoga* sp. TBGT1765 (1), *Thermotoga* sp. Mc24 (1), *Pseudothermotoga elfii* NBRC107921 (1), *Pseudothermotoga elfii* lettingae (2) |
| **T1074V1** | *Thermosipho affectus* B11070 (1), *Thermosipho affectus* 1223 (3),<br>*Thermosipho affectus* B11063 (1) |

\*Provirus that have been induced and shown to produce virus particles.

### Classification, genomic features and distribution of Group 2 proviruses

Group 2 consists of three complete proviruses in the genomes of *Petrotoga* sp. 8T1HF07 (P8T1HF07V1), *Petrotoga olearia* (POV1) and *Defluviitoga tunesiensis* (DTV1) (**Fig. 2**), and two incomplete proviruses in the genomes of *Petrotoga mobilis* SJ95 and *Petrotoga* sp. 9T1HF07 (**Supplementary Fig. S5**). Due to the short length of the incomplete proviruses, they were not included in the remaining analyses of this section.

Following the taxonomic classifications criteria described above, the three complete proviruses P8T1HF07V1, DTV1 and POV1 are assigned to three new viral species (**Supplementary Table S4**). Based on the head-neck-tail module classification [33], these proviruses likely encode siphoviruses of Type1 - Cluster 2. All hosts of the previously described members of this *Siphoviridae* lineage belong to Firmicutes. In agreement with this, similarity searches revealed that these proviruses show very high similarity to proviruses of three Firmicutes genomes: *Lacticigenium naphta*e (LNV1), *Geosporobacter ferrireducens* (GFV1) and *Mahella australiensis* (MAV1) (**Fig. 2, Supplementary Table S4**).

The sequences and genome organization of the three complete Group 2 proviruses differ considerably from that of Group 1 (**Fig. 1** and **Fig. 2**). These proviruses are also not located next to tRNA genes. The 5’ module encodes genes involved in virus replication and transcription, and the comparative genomic analysis shows high level of diversity in this region (**Fig. 2**). This module is followed by highly conserved packaging, morphogenesis and lysis modules. The lysogeny module is located at the 3’ end of the virus. The site-specific serine recombinases carried by the Group 2 proviruses in this module are distant homologs of the earlier discussed Group 1 recombinases (**Fig. 4**).

When comparing the Group 2 provirus genomes from Thermotogota and Firmicutes, each Thermotogota provirus is more similar to a Firmicutes provirus than to other Thermotogota proviruses **(Fig. 2, Supplemental Table S4).** Alignments of the two Thermotogota-Firmicutes provirus pairs, P8T1HF07V1 and GFV1, and DTV1 and MAV1, have 66.7 % and 56.7 % intergenomic similarity values, respectively (**Supplementary Table S4**), suggesting they may be assigned to the same genus. Moreover, P8T1HF07V1 has 97% nucleotide identity to the GFV1 over specific subregions that encode structural genes and the genes for DNA packaging and genome integration **(Fig. 3)**. Similarly, in the same regions in DTV1 and MAV1 have 95-97% identity. In contrast, the same regions in P8T1HF07V1 and POV1, and P8T1HF07V1 and DTV1 have 53 and 75% nucleotide identity, respectively. Such similarity patterns suggest that these viruses likely can infect hosts from both Thermotogota and Firmicutes phyla.

In contrast to the Group 1, none of the Group 2 proviruses had matches to CRISPR spacers in 90 Thermotogota genomes, suggesting that the Group 2 viruses have a more restricted host range within the Thermotogota or started to infect members of this phylum recently.

### Successful induction of TAV1 from *T. africanus* H17ap60333

Induction assays were performed on three of the putatively lysogenized *Thermotogota*: *T. africanus* H17ap6033 (Group 1), *Petrotoga* sp. P8T1HF07 (Group 2) and *P. olearia* (Group 2). Only the provirus in *T. africanus* H17ap6033 (TAV1) was successfully induced using mitomycin C. TAV1 was shown to produce viral particles with a polyhedral head of ~50 nm in diameter and a flexible non-contractile tail of ~160 nm in length and ~10 nm in width (**Fig. 5a**). Based on tail morphology, TAV1 was classified to the order *Caudovirales* and the family *Siphoviridae*, confirming the sequence-based classification. TAV1 morphology is similar to the three previously characterized temperate *Marinitoga* viruses, whose virion tails were just slightly longer [15, 16]. In addition to viral particles, a release of membrane vesicles or toga fragments was regularly observed (**Fig. 5b**).

**Fig. 5.**
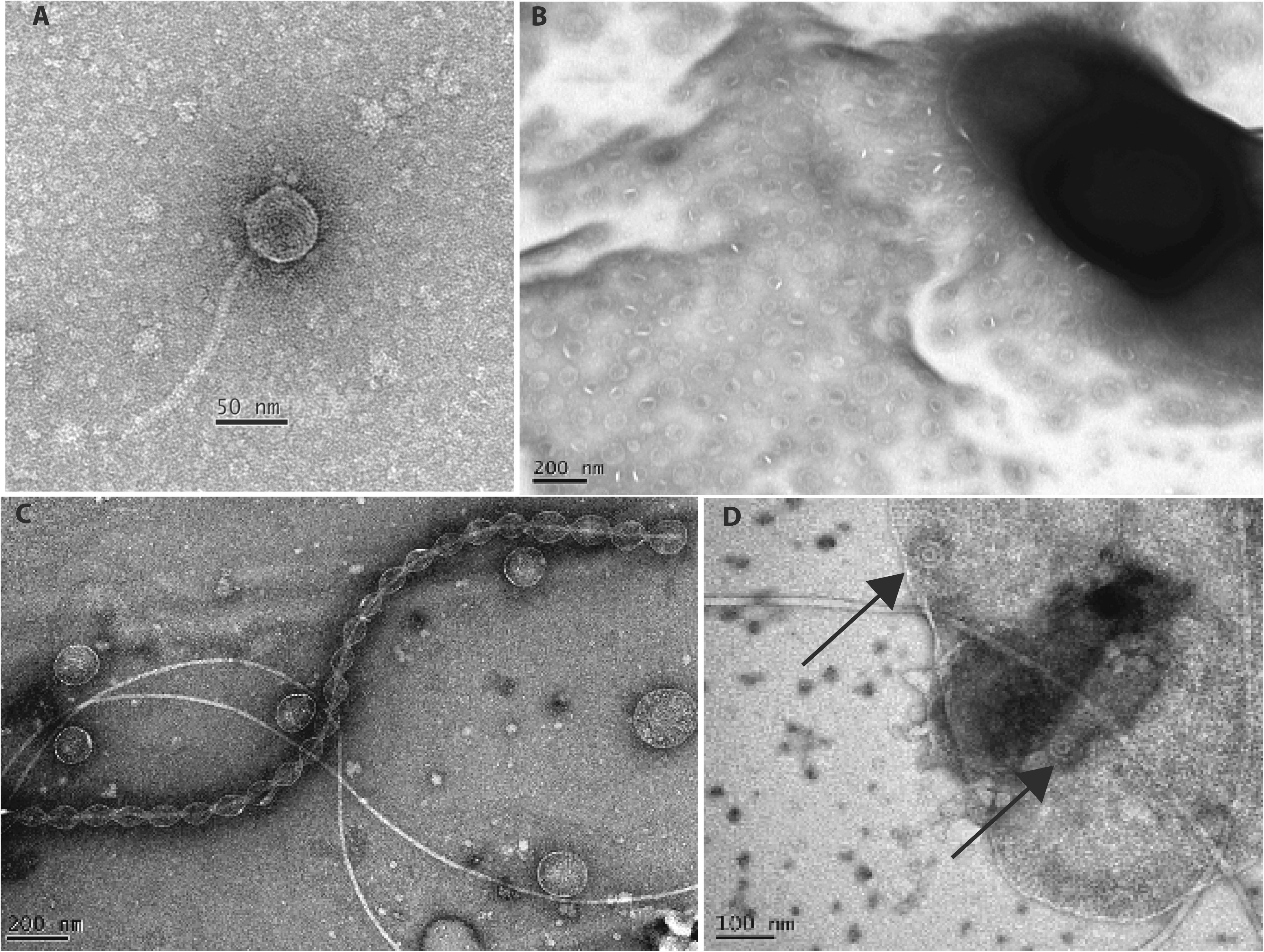
Electron micrographs of the induced virus and vesicles, stained with 2% uranyl acetate. **Panel a.** The TAV1 virus particle, which shows a typical Siphoviridae morphology. **Panel b.** Vesicles and toga fragments produced by *Thermosipho africanus* H17ap60334. **Panel c.** Vesicles produced by *Petrotoga* sp. 8T1HF07.NaAc.6.1, some of which are attached to a flagellum. **Panel d.** Sunflower-like structures inside *Petrotoga* sp. 8T1HF07.NaAc.6.1 cells. The structures are highlighted by arrows.

While the induction of the proviruses in *Petrotoga* sp. 8T1HF07 (P8T1HF07V1) and *P. olearia* (POV1) using mitomycin C was unsuccessful, membrane vesicles of various sizes and shapes (20 - 100nm) were produced by the cells, and in particular by the induced *Petrotoga* sp. 8T1HF07 cells. Analysis of the supernatant of the latter culture revealed similarly-sized round-shaped vesicles connected together in long chains by hooking onto the flagella, like a “pearl necklace”, while free vesicles showed more diversity in size and shape (**Fig. 5c**). Some “sunflower-like” structures were also observed inside a remaining cell (**Fig. 5d**). It is unknown if the provirus or stressors influence the production of these vesicles and structures, or if they are produced spontaneously.

### A potential receptor for the Group 2 viruses

Several types of structures on the surface of bacteria, such as membrane proteins, flagella, pili, or carbohydrate moieties, can act as virus receptors [47]. Most siphoviruses of Gram-negative bacteria, and some of Gram-positive bacteria, use proteinaceous receptors for adsorption [48, 49]. If the Group 2 viruses use the same protein receptor to attach to both Thermotogota and Firmicutes cells, the large phylogenetic distance between these hosts offers an opportunity to identify possible membrane protein receptors bioinformatically, since the receptor proteins would be expected to be conserved across the genomes from both phyla. It should be noted that this approach would only identify possible protein receptors, while potential shared carbohydrate receptors would not be detected.

Four predicted membrane proteins with transmembrane helices were identified in all genomes carrying a Group 2 provirus. One of these was the viral holin gene, leaving three receptor candidates: a ComEA family DNA-binding protein, an oxaloacetate decarboxylase beta subunit, and an ABC transporter ATP-binding protein. Phylogenetic analyses revealed that the ComEA and the oxaloacetate decarboxylase homologs are widely distributed among Thermotogota (**Supplementary Fig. S6**). In contrast, the ABC transporter is, among the Thermotogota, restricted to *Petrotoga* and *Defluviitoga*, the two genera where the Group 2 proviruses are observed (**Supplementary Fig. S6, panel C**).

Moreover, the phylogenetic analysis suggests the homologs in *Petrotoga* and *Defluviitoga* originated from an LGT event with a Firmicute (**Supplementary Fig. S6)**. These proteins show particularly high amino acid sequence similarity in the C-terminal domain of both Thermotogota and Firmicutes homologs, which is facing the exterior of the cell and could serve as a virus target (**Supplementary Fig. S7**). Although experiments are needed to demonstrate if any of these proteins functions as receptor for these viruses, we suggest that the ABC-transporter ATP-binding protein is a strong candidate for a Group 2 virus receptor.

### Moron genes are abundant in the identified proviruses

Many temperate viruses are known to carry moron genes, which are genes that do not have a direct viral function [50, 51]. The detected proviruses of Thermotogota are no exception: the Group 1 proviruses carry up to 6 morons (**Fig. 1**), while the Group 2 proviruses have between 4 and 13 morons (**Fig. 2**). However, it should be noted that because the 5’ ends were hard to define for Group 2 proviruses, some of the moron genes at the 5’ ends might not be part of the proviruses. Sequencing virus DNA isolated from capsids will help resolve this issue in the future.

Among the morons are several proteins that may confer a selective advantage to the host (**Fig. 1**and **Fig. 2**, **Supplementary Table S2**). For instance, M1138V1 carry two genes involved in sulfur metabolism. The Group 2 proviruses encode several transporters, peptidases and hydrolases, likely to be beneficial for these heterotrophic bacteria. In addition, all the viruses carry several hypothetical proteins that may also have non-viral functions.

### Evidence for the viruses’ impact on lateral gene transfer

Eight hundred seventy homologs of 106 proviral genes were detected in 54 out of 59 Thermotogota genomes with no detectable proviruses (**Supplementary Table S6**). It should be noted that some provirus genes, e.g. the Group 1 recombinases and terminases (**Fig. 4** and Supplementary **Fig. S3**), did not pass our stringent screening criteria (see Material and Methods), thus these represent minimum estimates of matches to proviral genes in these genomes. Notably, 370 of 870 were homologs of 28 moron genes, suggesting that the viruses may facilitate exchange of “host” genes among Thermotogota. Moron genes also had the highest number of homologs across 54 genomes, with most abundant being a queuine tRNA-ribosyltransferase in the Group 2 provirus DTV1 (found in 48 genomes) and an aldo/keto reductase in Group 1 provirus GPV1 (found in 41 genomes).

Among phylogenetically informative datasets, 10 proviral genes group within Thermotogota and 17 group within Firmicutes, suggesting that many of the proviral genes originated either in Thermotogota and Firmicutes (**Supplementary Table S6)**. For instance, the above-described abundant moron gene queuine tRNA-ribosyltransferase is of Thermotogota origin, while the aldo/keto reductase appears to be of Firmicutes origin (**Supplementary Table S6)**. In the phylogeny of another moron gene, a cadmium or heavy metal transporter found in Firmicutes provirus MAV1 and Thermotogota Group 2 proviruses DTV1 and POV1, the provirus genes group closely with Firmicutes’ homologs (**Supplementary Fig. S8**). Notably, the other closely related Thermotogota homologs are found in three *Fervidobacterium* and *Pseudothermotoga* genomes, genera where no proviruses have yet been identified. Inspecting the genomic region surrounding these genes in *Fervidobacterium* and *Pseudothermotoga*, revealed that the homolog of the proviral recombinase (**Fig. 4**) is located immediately upstream of the transporter gene. No other typical virus genes were observed in these regions, suggesting these genes are likely remnants of proviruses. This also indicates that viruses related to Group 2 proviruses may have broader host range that we presently detect.

Taken together the above analyses suggest that the viruses of both Group 1 and Group 2 may facilitate exchange of genes not only among Thermotogota, but also between Thermotogota and Firmicutes.

## Discussion

In our search for proviruses in genomes of Thermotogota, we discovered two distinct groups of temperate siphoviruses that have lysogenized this bacterial phylum. These proviruses may represent multiple new viral species and genera. Our analyses suggest that these viruses likely have broad host range that spans at least multiple genera. We also found that the identified proviruses lineages are closely related to Firmicutes’ viruses.

One of the bioinformatically identified Group 1 proviruses (TAV1) was induced and shown to produce virus particles. The provirus resides in a genome of a *T. africanus* isolate from the Hibernia oil reservoir off the Canadian east coast. The analysis of CRISPR spacers suggested that this virus may have a particularly wide host range, with the highest number of spacer-matches in genomes from outside its genus. For instance, a virus very similar to TAV1 had likely infected *Thermotoga* spp. isolates from the North Sea Troll oil reservoir. Similar predatory virus pressure in geographically and geologically remote subsurface environments have been observed for *Methanohalophilus* isolates from reservoirs in the USA and Russia [52].

We were not able to induce virus production from the selected Group 2 proviruses This could be due to these proviruses currently being inactive, or we may not have applied the right conditions to induce the expression of these proviruses. Nevertheless, the high level of sequence identity between Group 2 provirus sequences from Thermotogota and Firmicutes phyla uggests that they have been active very recently.

Many of the genes carried by both Group 1 and Group 2 proviruses are found in genomes of Thermotogota that do not have detectable proviruses. These genes often group with Firmicutes or viruses that infect Firmicutes, and many can be classified as morons. This suggests that both Group 1 and Group 2 viruses transfer genes within Thermotogota and between Thermotogota and Firmicutes, and may serve as a major mechanism for the earlier reported large amounts of lateral gene transfer between Thermotogota and Firmicutes [7, 8].

Based on our bioinformatic and phylogenetic analyses, we propose that an ABC transporter may serve as a receptor for at least some of these proviruses. ABC transporter proteins are, to our knowledge, not commonly identified as bacterioviral receptors. However, *Lactoccocus* viruses from the siphoviral c2 group have been shown to use membrane proteins Pip or YjaE, both with sequence similarity to ABC-transporter domains, as secondary receptors [47, 53].

Intriguingly, transporters, and ABC transporters in particular, are among the most frequently transferred genes both within the Thermotogota and between Thermotogota and Firmicutes [7, 8, 54]. Transporter genes were also detected in the provirus genomes. The possibility that transporters can function as viral receptors in the Thermotogota therefore suggests that acquiring a new transporter, perhaps via a viral infection, might result in the cell not only acquiring a new function but also becoming susceptible to a new virus. This virus might carry another transporter gene, which can introduce yet another virus, resulting in a ratchet-like process. Using transporters as receptors will therefore not only provide the virus with the wide host range but could also make viruses the vehicles on the highways of gene sharing observed between the Thermotogota and Firmicutes.

Genes encoding proteins for membrane transport, including ABC transporters, have been observed in several other viruses [55], and thus the proposed process could operate widely among bacteria. This is contrary to a role commonly assigned to morons where they often confer resistance to infections by other viruses [51]. Further studies and experiments are needed to investigate if such ratchet processes are indeed occurring in natural systems. However, regardless of the functions of the morons in the *Thermotogota* proviruses, the observation of viruses potentially infecting organisms from different phyla further demonstrates that viruses are key actors in the evolution of microbial diversity.

## Acknowledgements

This work is supported by a Research Council of Norway award (project no. 180444/V40) to C.L.N., by the Sino-French LIA/PRC 1211 MicrobSea to J.L. and by the Simons Foundation Investigator in Mathematical Modeling of Living Systems award 327936 to O.Z. Strains were obtained from the Université de Bretagne Occidentale Culture Collection (UBOCC, Plouzané, France, www.univ-brest.fr/ubocc).

## Supplementary Figures

**Supplementary Fig. S1. Panel A. Gene-sharing network of proviruses calculated in VContact2.** The network is based on shared protein clusters between viral genomes. Only proviruses at most three nodes away from MPV1 and P8T1HF07V1 are shown. The Thermotogota proviruses are colored in red, viruses from Firmicutes are blue and viruses infecting other taxa are colored orange. The quality scores calculated by ClusterOne are 0.94 (p=0.00004) for the Group 1 cluster and 0.83 (p=0.006) for the Group-2 cluster. **Panel B. Placement of proviruses on the phylogenetic tree of Thermotogota genomes reconstructed from 74 single copy protein-coding genes**. Closely related genomes (distance > 0.1), monophyletic genomes from the same genus, and clades consisting of only metagenome assembled genomes were collapsed. Identified proviruses are indicated next to their respective host genera. The tree was visualized in iTOL [56]. Tree scale, substitutions per site.

**Supplementary Fig. S2**. **Placement of complete Thermotogota and Firmicutes proviruses on the viral proteomic tree.** The viral proteomic tree is from ViPTree v. 1.9 [35], and only the relevant region of the tree is shown. The Thermotogota and Firmicutes proviruses are labeled with red stars. Taxonomy of the related viruses and their hosts is indicated as color bars next to a terminal leaf on the tree.

**Supplementary Fig. S3. Maximum likelihood trees of three families of terminase large subunit genes.** The phylogenetic trees displayed were constructed using RAxML as implemented in Geneious v. 10 with a GAMMA-WAG substitution model and 100 bootstrap replicates. The trees should be considered unrooted. Bootstrap support > 70% is shown on branches as circles, with the size corresponding to the strength of support. Taxonomic labels of Thermotogota with proviruses are shown in red bold font, with the provirus name listed after the host name. Thermotogota homologs from genomes with no detected provirus are listed in bold font. Numbers in front of each taxon name represent database accession numbers. The tree was visualized in iTOL and rooted by midpoint rooting and should be considered unrooted [56].

**Supplementary Fig. S4. Overview of CRISPR spacer sequences from Thermotoga isolates from the Troll oil reservoir mapped on to the TAV1 genome.** Alignment position of each CRISPR spacer is indicated as black bars. Mapping and visualization was performed in Geneious v. 10 and maximum of one mismatch was allowed.

**Supplementary Fig. S5. Comparison of the three complete and two incomplete Thermotogota Group 2 provirus sequences.** Virus name and the genus the host belongs to is indicated. The regions with significant pairwise BLASTX similarity scores are connected, red indicates that sequence is in the same direction while blue indicates that the similar sequences are on opposite strands. The predicted ORFs are color-coded based on their function and should be considered approximate, because it relies only on gene annotations. Selected gene annotations are included and abbreviated; HMT ATPase: heavy metal translocating ATPase, FMN reductase: flavine mono nucleotide reductase, HAD family phosphatase: haloacid dehalogenase superfamily of hydrolase), dimethyladenosine trf: dimethyladenosine transferase, 2Fe-2S bp: 2Fe-2S binding p rotein, MFS transporter: multi facilitator superfamily transporter. The figure was produced using genoPlotR [27].

**Supplementary Fig. S6. Maximum likelihood trees of three potential virus receptor genes. Panel A: Competence protein ComEA, Panel B: oxaloacetate decarboxylase and Panel C: ATP-binding cassette, subfamily B.** Bootstrap support > 70% is shown on branches as circles, with the size corresponding to the strength of support. The names of Thermotogota taxa that contain Group 2 proviruses are displayed in red font and Firmicutes with Group 2-like proviruses are displayed in blue font. Clades containing sequences from the same genus are collapsed into wedges. The trees were rooted using midpoint rooting, and should be considered unrooted. The trees were visualized in iTOL [56].

**Supplementary Fig. S7. Overview of the alignment of the ABC transporter ATP-binding protein in Thermotogota and Firmicutes genomes with Group 2 proviruses.** Sites 100% conserved in all sequences sites are highlighted in color, while variable sites are shown in grey. Transmembrane regions, predicted using the TMHMM Server v. 2.0, are shown in red above the alignment (http://www.cbs.dtu.dk/services/TMHMM-2.0/).

**Supplementary Fig. S8. Maximum likelihood tree of the moron gene annotated as a cadmium transporter**. Bootstrap support > 70% is shown on branches as circles, with the size corresponding to the strength of support. Taxon names of Thermotogota with a provirus are given in red and taxon name of Firmicutes with a Group 2-like provirus are given in blue. Provirus name is also indicated. Thermotogota homologs from genomes with no detected provirus, or where the homolog is found outside the provirus region, are given in bold font. Database accession numbers are shown in front of taxonomic names. The tree was rooted by midpoint rooting and visualized in iTOL [56].

