## Supplementary figures and images for "Newly identified proviruses in Thermotogota suggest that viruses are the vehicles on the highways of interphylum gene sharing"

### Supplementary Fig. S1

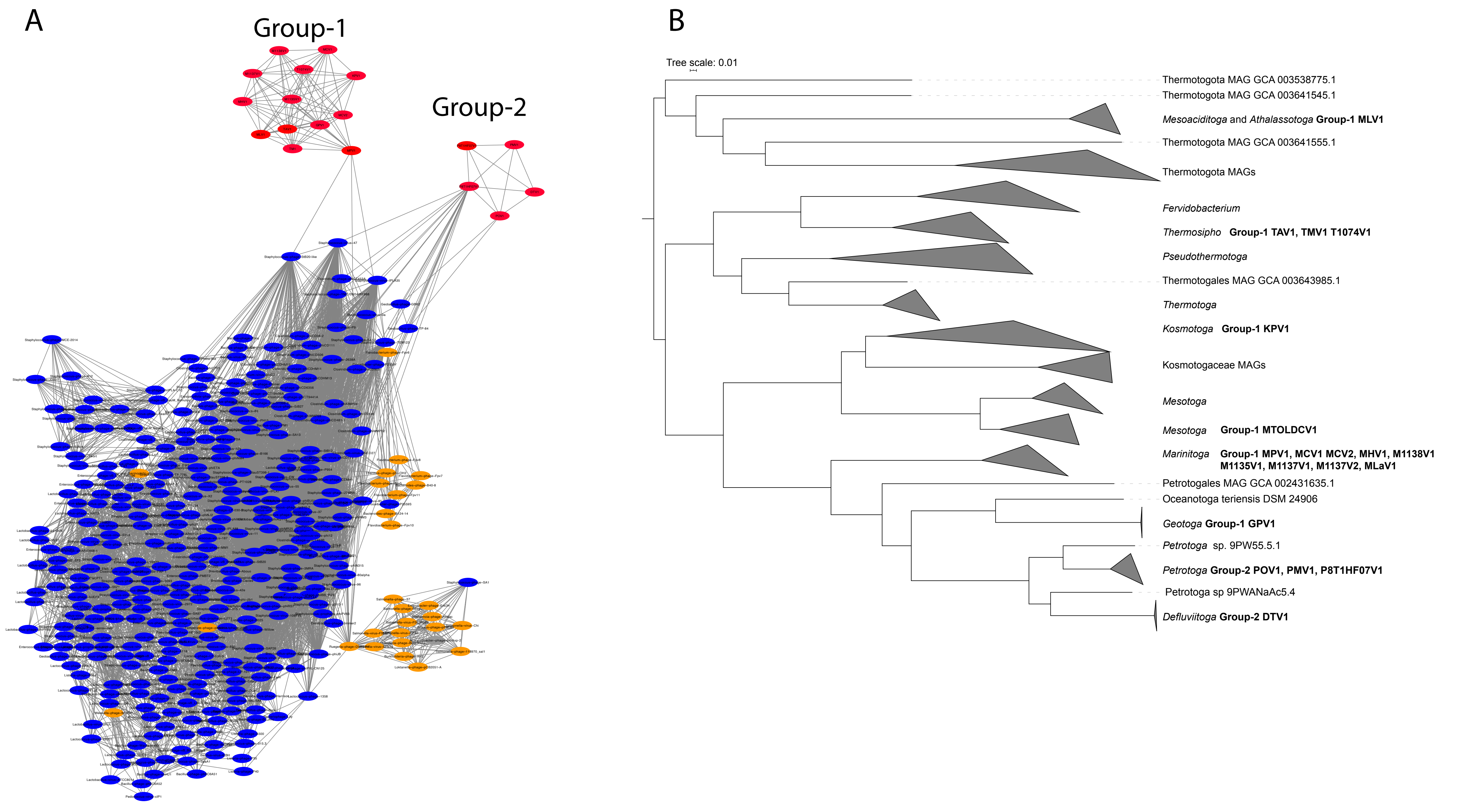

### Supplementary Fig. S2

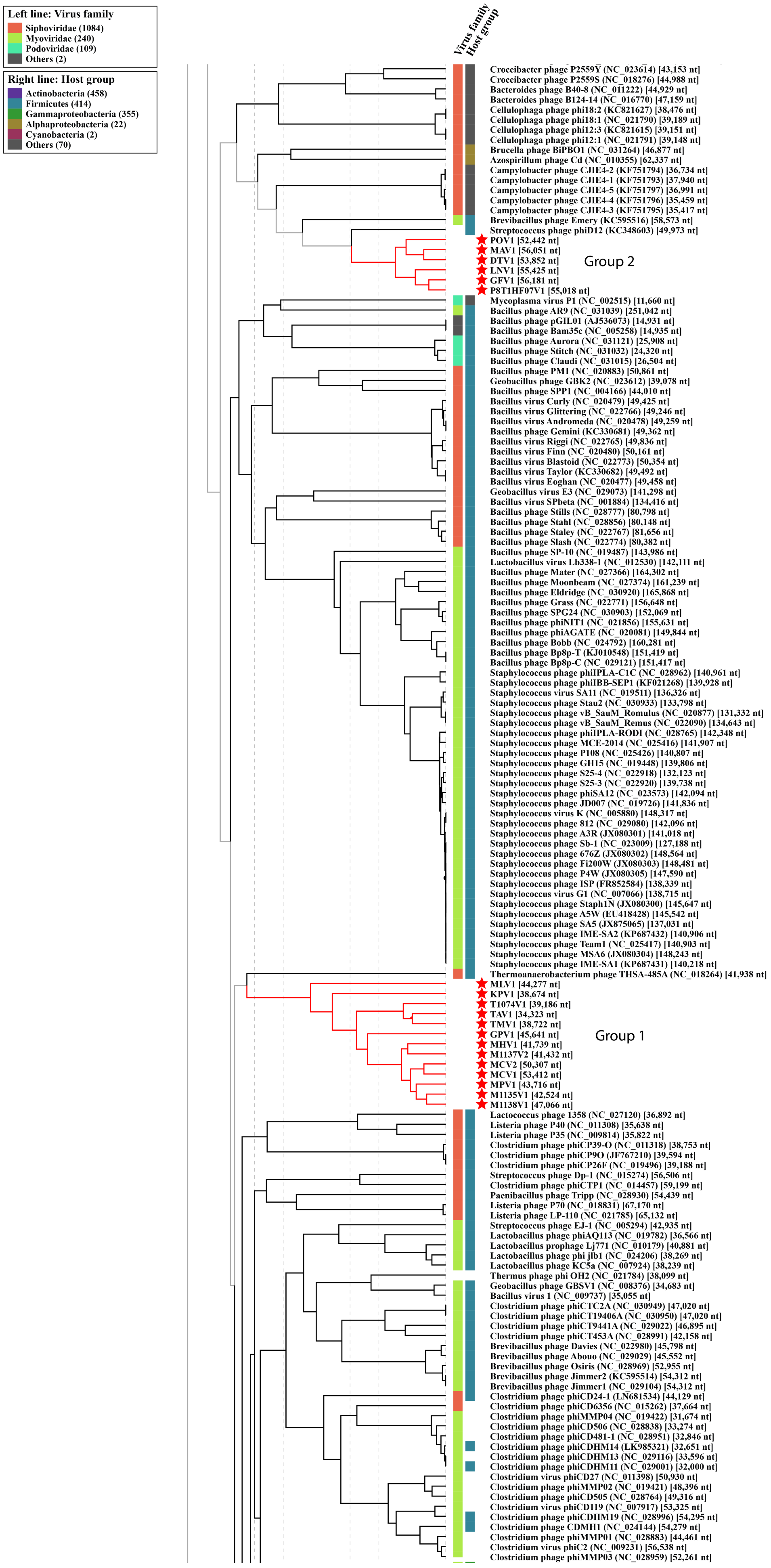

### Supplementary Fig. S3

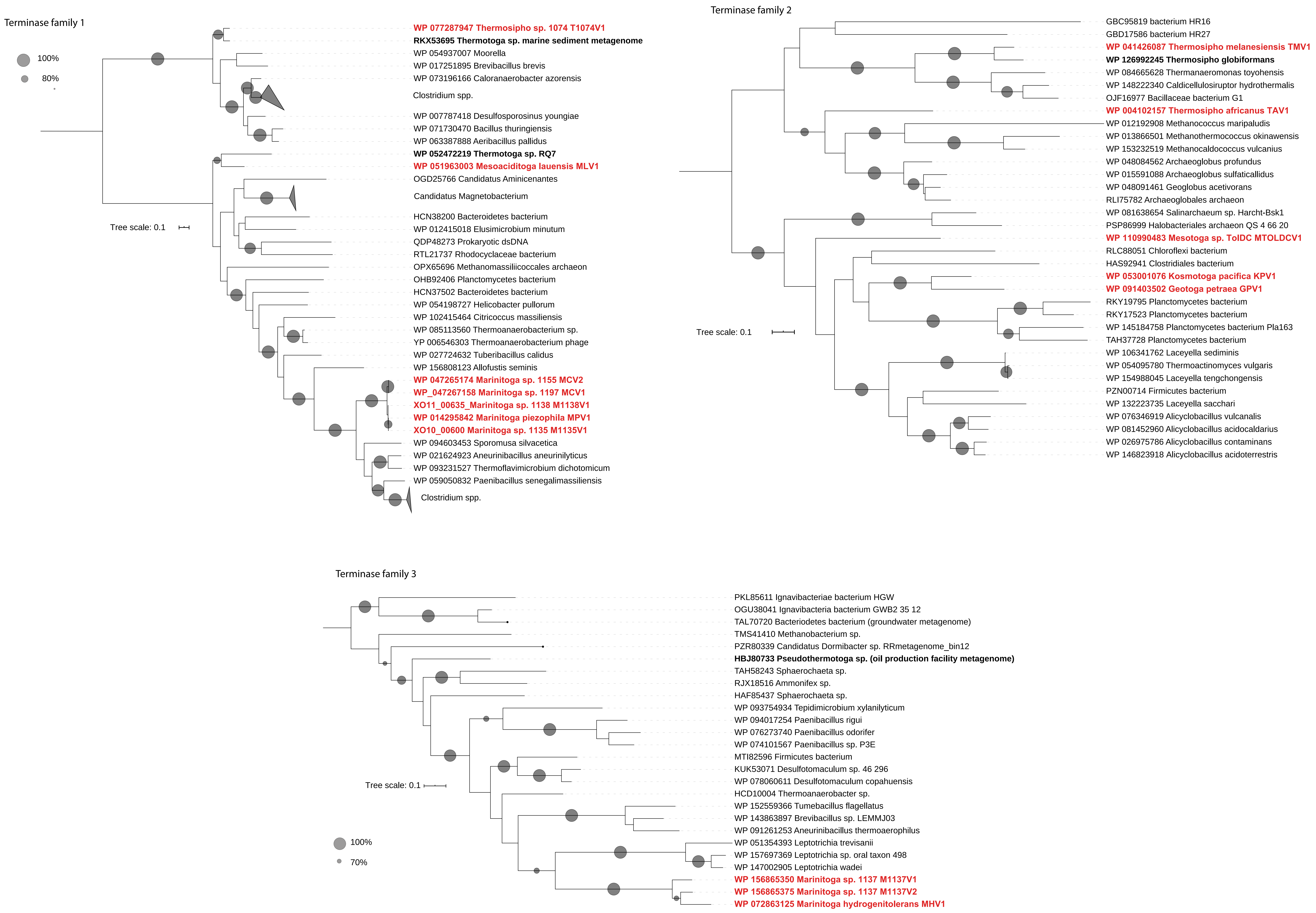

### Supplementary Fig. S4

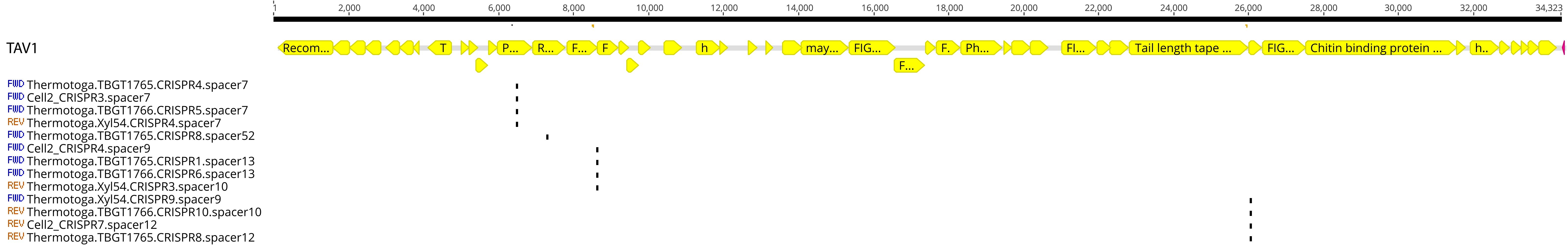

### Supplementary Fig. S5

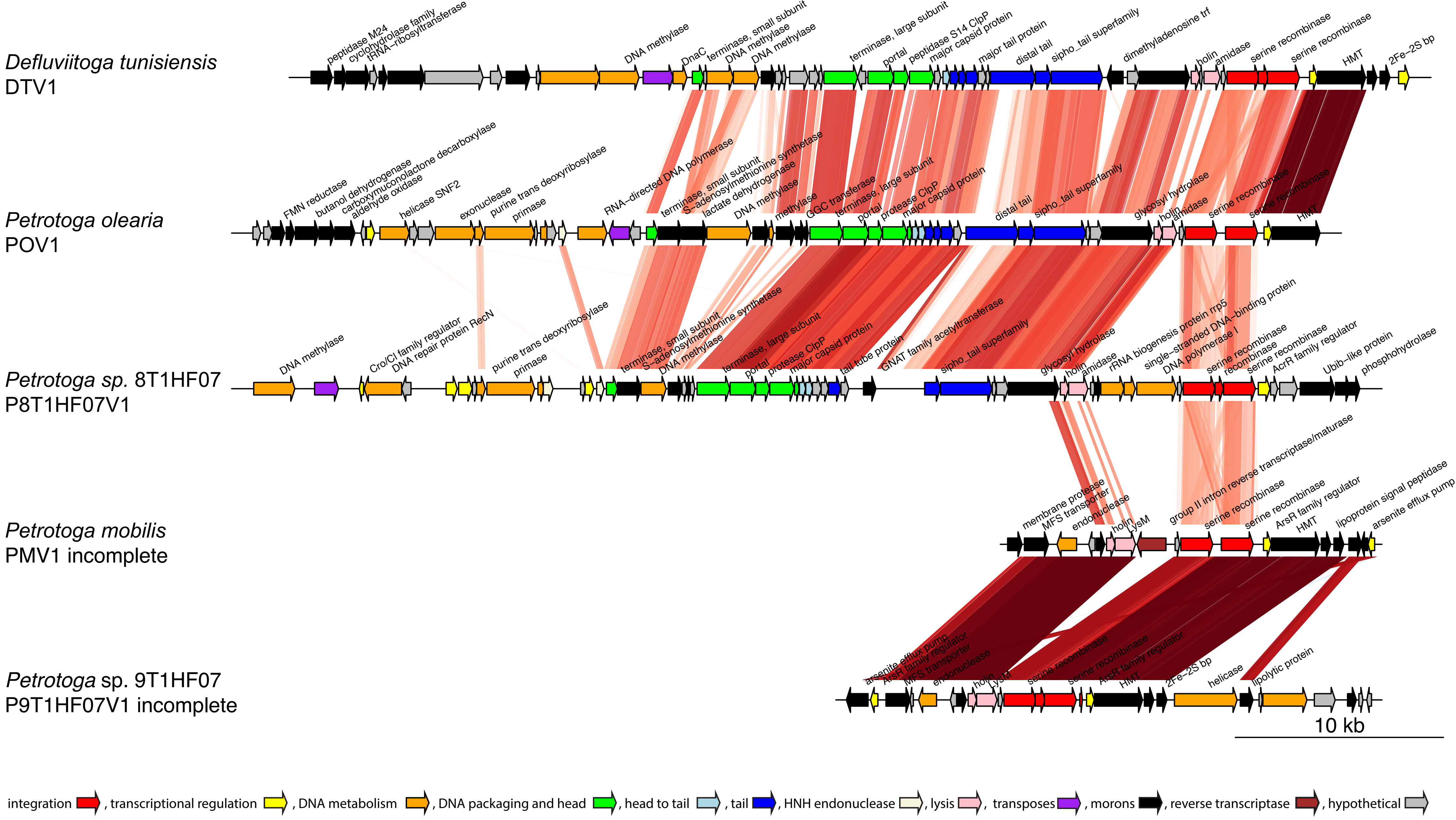

### Supplementary Fig. S6

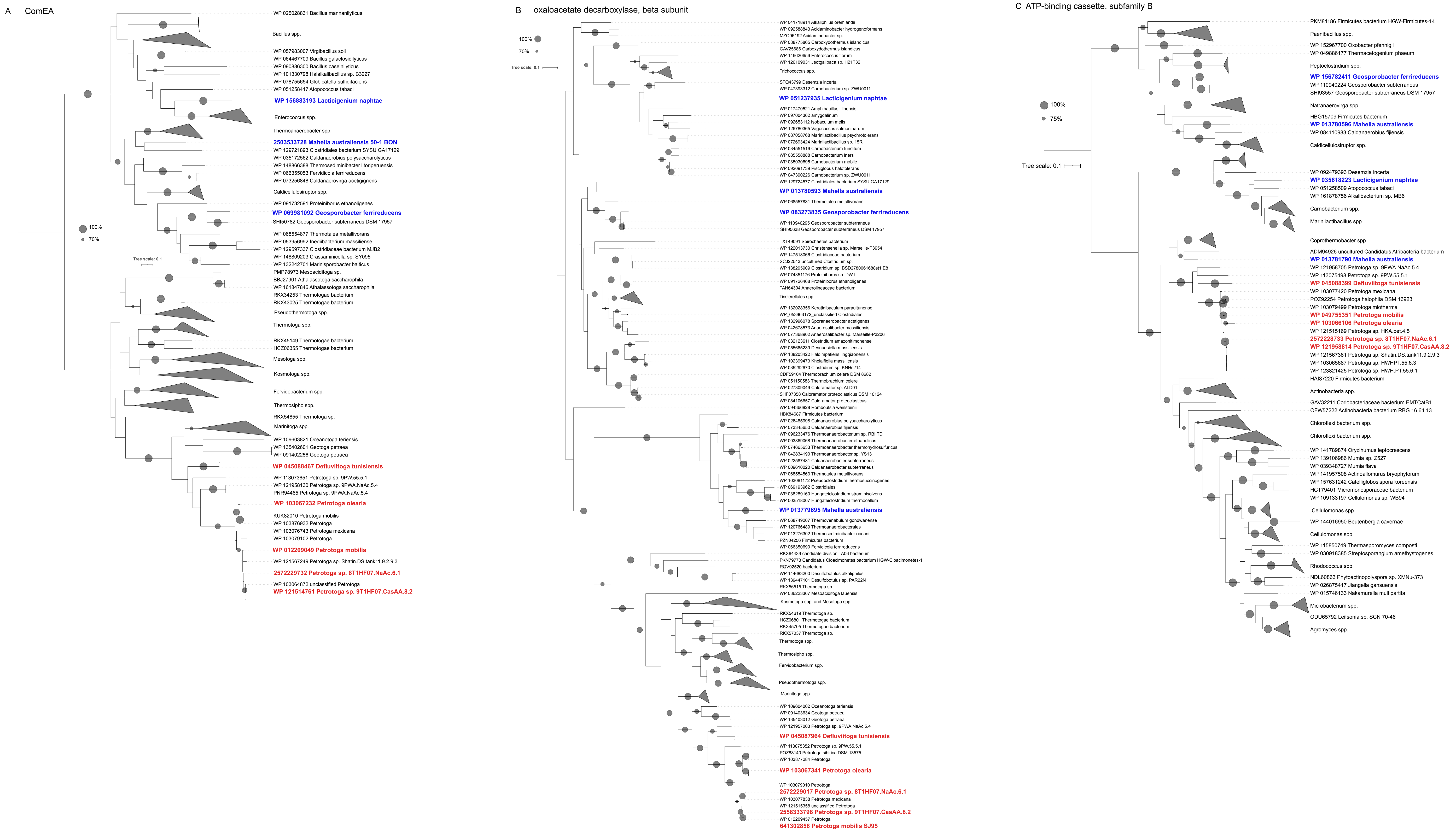

### Supplementary Fig. S7

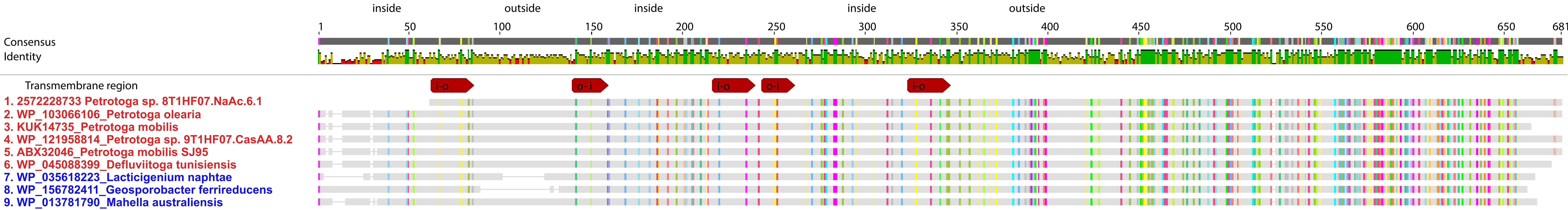

### Supplementary Fig. S8

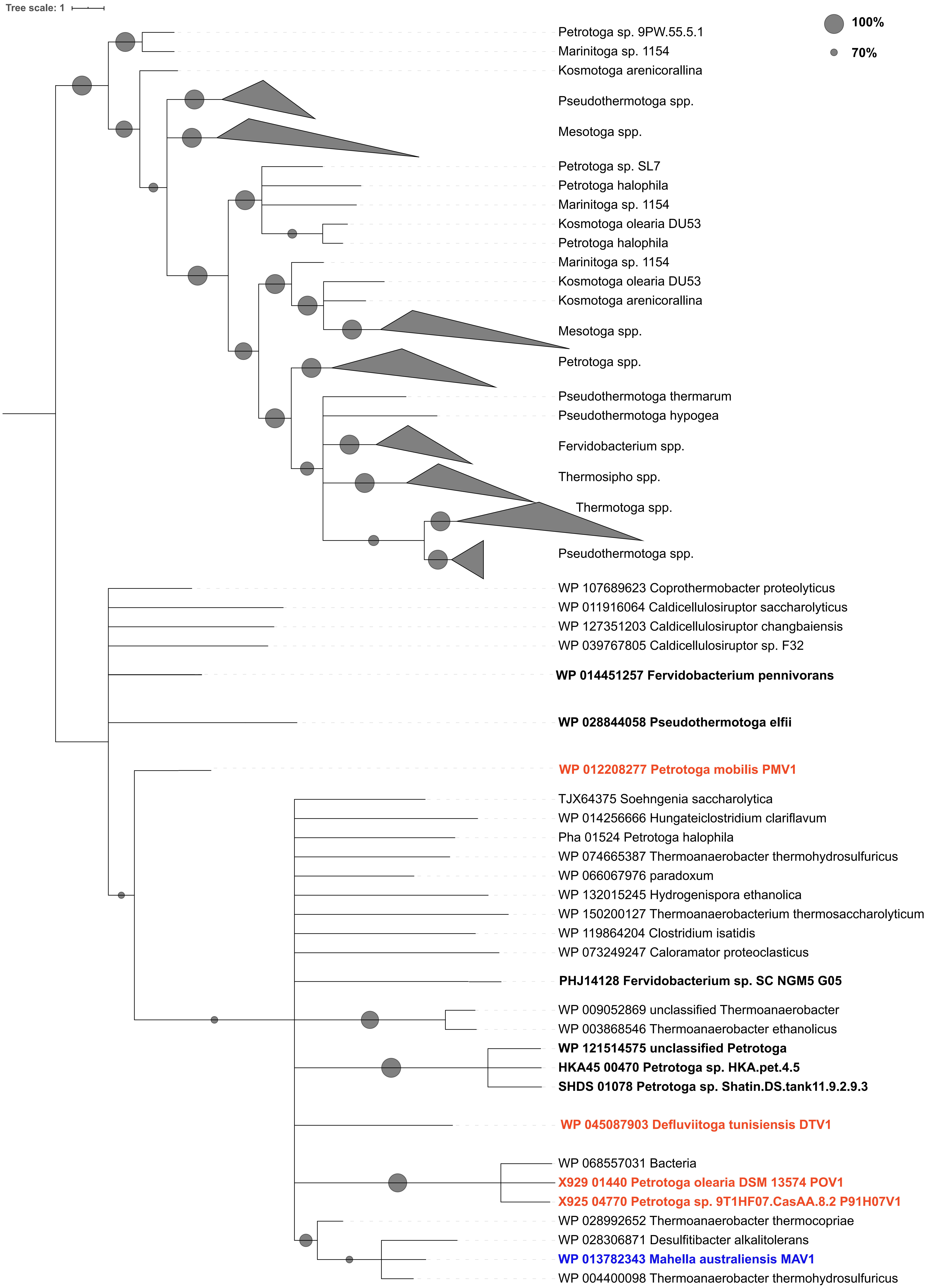
